# A non-catalytic function for Rad18 in sustaining glioblastoma proliferation

**DOI:** 10.1101/2024.09.05.611406

**Authors:** Chames Kermi, Nour Benbahouche, Lenka Stefancikova, Aurore Siegfried, Jean-Marc Pascussi, Julie Pannequin, Jérôme Moreaux, Tom Egger, Jean-Philippe Hugnot, Marie-Bernadette Delisle, Elizabeth Moyal, Emmanuelle Uro-Coste, Domenico Maiorano

**Author notes:** Corresponding author Contact information: Institut de Génétique Humaine. UMR9002 CNRS-UM 141, rue de la Cardonille. 34396 Cedex 5 Montpellier. France. « The authors declare no potential conflicts of interest ».

## Abstract

The Rad18 E3 ubiquitin ligase, a non-essential gene, is a key regulator of DNA damage tolerance that also functions in repair of DNA double strand breaks. Rad18 is overexpressed in the aggressive brain cancer glioblastoma (GBM) and its downregulation sensitizes glioblastoma cells to DNA damaging agents. Here we show that Rad18 has an essential role in GBM cells proliferation in the absence of external damage, surprisingly independent of its catalytic activity. Rad18 downregulation leads to cell cycle arrest in the G1 phase in the absence of apparent DNA damage. We also show that Rad18 sustains GBM stem cells self-renewal and survival, as well as the growth of tumor orthotropic xenografts in mice. We also show that increased Rad18 expression enhances the growth of non-transformed cells and induces features of oncogenic transformation. Mechanistically, we show that Rad18 downregulation negatively regulates the Hippo pathway by interfering with the nuclear retention of the YAP1 transcription factor. Altogether, these data show that Rad18 has an essential, non-catalytic function, in GBM proliferation, and propose Rad18 as a key target to sensitize GBM to therapy.

## Introduction

Glioblastoma (GBM, World Health Organization grade IV glioma) is the most frequent and aggressive primary brain tumor. The mean survival time of patients affected by this disease is less than two years due to a strong resistance to standard-of-care therapy, consisting of surgical resection followed by radiotherapy and chemotherapeutic treatment with alkylating agents (temozolomide), whose molecular bases are currently poorly understood (1). GBM stem cells (GSCs), isolated from samples of human glioblastoma, are currently considered as best candidates to explain glioblastoma aggressiveness and resistance to the therapy (2). GSCs express pluripotency markers (Oct4, Sox2, Nanog amongst others), are characterized by self-renewing, multipotency (ability to generate differentiated cells of various types) and tumorogenicity. Amongst various cellular pathways controlling the expression of the pluripotency factors such as Sox2, is the YAP-Hippo pathway (3). When transplanted into immunodeficient mice GSCs are able to form tumors that closely resemble the initial human pathology. GSCs have the particularity to form floating clonal colonies growing in suspensions, also known as gliomaspheres. Being stem cells, they also have the ability to generate the three cell lineages of the brain upon differentiation, such as neurons, astrocytes and oligodendrocytes, which display a marked decrease in their tumorigenic potential. Targeting genes specifically involved in GSCs growth and maintenance therefore constitutes a promising approach in the treatment of glioblastoma (4).

We have previously reported that the non-essential Rad18 E3 ubiquitin ligase, a critical regulator of DNA damage tolerance involving translesion DNA synthesis (TLS), and also involved in DNA double strand break (DSBs) repair (5) for review), is abundant in GSCs compared to their differentiated counterparts (6), suggesting a potential function in GSCs biology. Consistent with this observation, few recent reports have linked high Rad18 expression with GBM survival and prognosis (7–9), although the molecular grounds of this effect have not been explored. Following our key observation (6), we have explored the significance of increased Rad18 expression in GBM and found that Rad18 is by itself required for proliferation of adherent GBM cells in culture, independently of its catalytic activity. We have also observed that Rad18 downregulation arrest cells in the G1 phase of the cell cycle, without detectable DNA damage. We further show that Rad18 is important for GSCs growth in culture and in mice xenografts in the absence of external damage, and that Rad18 has an oncogenic potential by regulating the Hyppo pathway.

## Material & Methods

### Cell culture

U87 (RRID:CVCL_GP63), LN18 (RRID:CVCL_0392), U251 (RRID:CVCL_0021), T98G (RRID:CVCL_0556) and NIH3T3 (RRID:CVCL_0594), cells were maintained in Dulbecco’s modified eagle’s medium (DMEM) supplemented with 10 % fetal bovine serum, 2 mM glutamine and antibiotics in a humidified atmosphere of 5 % CO_2_ at 37 °C. GSCs (Gb4 and Gb7) were isolated from fresh glioblastoma primary tumors and grown and maintained as previously described (10). Gb4 and Gb7 were fully characterized by CHG array. Gb4: EGFR, cMET, BRAF amplification, partial loss of chromosome 9p, 16p and 17q (NF1 loss); Gb7: EGFR, cMET BRAF, CDK4, MDM2, Tp53, NF1 amplification, CDK2NA and PTEN deletion. For transient expression of Rad18 or empty vector (pcDNA3; RRID:Addgene_15475), HEK293T cells (RRID:CVCL_HA71) were transfected using calcium phosphate. Cells were collected at indicated time points after treatment and rinsed once in PBS. Whole cell extracts were clarified by centrifugation at 12, 000 *g* for 10 min at 4 °C. Protein concentration of the clarified lysates was estimated using BCA method (Pierce). Equal amounts of proteins were used for western blot analysis.

### Generation of stable U87 and GSCs cells expressing inducible shRNAs

Lentiviral particles expressing either doxycycline-inducible control or Rad18-targeting shRNAs (Dharmacon, see below) were added to cells at 3 MOI per cells and incubated at 37 °C for 1 hour. Cells were then transferred into flasks and after a 24 hours incubation at 37 °C, cells were selected with puromycin (2 µg/mL) for 48 hours at 37 °C. Subsequently cells were grown in puromycin-free medium. Induction of the shRNA was obtained by adding doxycycline (Sigma, 4 µg/mL) to the culture medium, and replaced every 48 hours.

### Cell proliferation experiments

For rescue experiments with either wild-type or mutated Rad18 variants, T98G cells were transfected with the indicated siRNA (30 pmoles) using the Lipofectamine™ RNAimax reagent (Thermofisher Scientific) and incubated at 37 °C for 3 days. At day 4, cells were transfected with 2 µg of each plasmid DNA previously described (6, 11) using jetprime (Polyplus) reagent and incubated at 37 °C for two more days. Cells were then collected and frozen in liquid nitrogen for further analysis.

### Generation of stable NIH3T3 cells expressing Rad18

NIH-3T3 and Platinum-E (Cell Biolabs) cells were grown in Dulbecco’s modified eagle’s medium supplemented with 10 % fetal bovine serum, 2 mM glutamine and antibiotics. For infection, viral particles were generated by transfecting Platinum-E ecotropic packaging cell line with retroviral vectors (pLPC-puro; RRID:Addgene_19835) encoding *Xenopus* Rad18 using Lipofectamine^®^ reagent (Invitrogen). The viral supernatant was collected and NIH3T3 cells infected. Forty-eight hours after infection, cells were selected in puromycin (2.5 μg/ml, Sigma)-containing medium. Selected populations were expanded and promptly used for experiments.

### Clonogenic assay

Exponentially growing NIH 3T3 cells (expressing either empty vector or RAD18) were seeded into 10 cm Petri dishes (1.2 x 10^5^ cells/dish). Cells were detached by trypsin and plated into 10 cm Petri dishes at a density of 100 surviving cells per dish and to 6-well plates in density of 50 surviving cells/well. After 9 days, the colonies were fixed and stained with 0.5 % crystal violet in 50 % methanol. The colonies were counted to determine the plating efficiencies and survival fractions.

### Antibodies

The following antibodies were used: Rad18 (Abcam, ab 79763; RRID:AB_1603946); β-Actin (Sigma, A1978; RRID:AB_476692); USP7 (Abcam, ab108931; RRID:AB_10862844); Histone H3 (Abcam, ab1791; RRID:AB_302613); GAPDH (abcam, ab9484; RRID:AB_307274); MCM2 (ab4461, Abcam; RRID:AB_304470); Sox2 (Thermo Fisher A301-739A; RRID:AB_1211354); Oct-4 (ab19857, Abcam; RRID:AB_445175); H2AX (Abcam, ab11175; RRID:AB_10694556); γH2AX (Cell Signaling, 2577; RRID:AB_2118009); p53 (clone DO-7, Dako; RRID:AB_2889978); p53, phospho-serine 15 (PS15, 9286, Cell Signaling; RRID: AB_331464); Rad6 (A300-281A, Bethyl; RRID: AB_263400); p21 (clone F-5, sc-6246, Santa Cruz Biotechnology; RRID: AB_628073); αtubulin (T6199, Sigma; RRID:AB_477583); Chk1 (Santa Cruz, sc-8408; RRID:AB_627257); Phospho-Chk1^ser345^ (Cell Signaling, 2341; RRID:AB_331212); YAP1 (13584-1-AP, Proteintech; RRID: AB_2218915); YES1 (20243-1-AP, Proteintech; RRID:AB_10697656); PPP1CA (67070-1-Ig, Proteintech; RRID:AB_2882379); PPP1CB (55136-1-AP, Proteintech; RRID:AB_10837232); PPP1CC (11082-1-AP, Proteintech; RRID: AB_2168085); YWHAE (66946-1-Ig, Proteintech; RRID:AB_2882270); YWHAZ (14881-1-AP, Proteintech; RRID:AB_2218248); YWHAH (15222-1-AP, Proteintech; RRID: AB_10863119).

### Immunofluorescence

Cells were spun on coverslips, washed in PBS and fixed in 4 % formaldehyde for 10 minutes at room temperature. After washing with PBS, cells were extracted with 0.5 % NP-40, 10 % sucrose in PBS for 5 minutes at room temperature. After washing with PBS, coverslips were saturated with 3 % BSA in PBS and incubated with primary antibodies over night in a humid chamber. After washing and incubation with secondary antibodies for 1 hour at room temperature, coverslips were mounted on slide with Prolongold antifade reagent containing DAPI (Invitrogen).

### si/shRNA

U87 cells were transfected either with siRNA Rad18 or siRNA Luciferase (Ctrl) as a control as previously described (6). Twenty-four hours after transfection, cells were trypsinized and seeded in 12 wells plates at a density of 10^4^ cells/well. For rescue experiments, T98G cells were reverse transfected either with an siRNA targeting the Rad18 3’UTR (5’-AAUUUCCUUGGGCAUUUAUAA-3’) or with siControl (Ctrl), using RNAimax (Invitrogen) at a density of 7.10^5^ cells/plate in a 60 mm plate. Seventy-two hours post transfection (Day 3), cells were counted and seeded at a density of 25 x 10^3^ cells/well in a 6 well plate. Twenty-four hours later (Day 4), cells were rescued with either 2 µg of empty vector (Ev), or RAD18 wt, RAD18 C28F, RAD18 C207F using JETPrime reagent (POLyplus) and cultured for twenty four hours. On day 5, cells were trypsinized, counted and harvested. GSCs were transfected with a previously described Rad18 siRNA (6) by electroporation (Nucleofector technology, Lonza) or the ShRAD18-3, 5’-TGGTTCTACATCAGAGTTGTT-3’ (Qiagen). Lentiviral particles expressing SMART vectors doxycyclin-inducible shRNA were from Dharmacon as follows: shRNA negative control (Non-targeting; ref. VSC6573); shRNA positive control (GAPDH; ref. VSH6545); shRNA targeting Rad18 (CAATCAAAGCTGGACTCCC, ref. V3IHSHER_8895752).

### Flow cytometry

U87 cells were transfected with the indicated siRNA as described above. Forty-eight hours post-transfection, cells were pulsed with BrdU for 15 minutes, harvested, washed twice in PBS and fixed in ice-cold 70% ethanol at -20 °C overnight. Thawed cells were washed twice in PBS and incubated with 50 µg/mL RNase A at 37 °C for 1hour. DNA was stained with propidium iodide (25 µg/mL). Cells were analyzed with a FACScalibur flow cytometer using CellQuestPro software.

### ELDA

After transfection with the indicated siRNA by electroporation, cells were diluted in pre-warmed growth medium, transferred to a flask containing 5 mL of medium and left to recover for 72 hours at 37 °C. Gliomaspheres were then dissociated and sorted by FACS in the presence of 7-AAD to distinguish live form dead cells. Then, a decreasing number of cells were seeded into 96-well plate pre-coated with Poly-HEME (Sigma, ref. P3932). Sphere formation was scored, up to seven days after seeding by phase contrast microscopy. Extreme limiting dilution analysis was performed using software available at bioinf.wehi.edu.au/software/elda/.

### Mice xenografts

All animal procedures were performed in accordance with a protocol approved by the local ethical committee. Glioblastoma-derived tumors were randomly assigned into control or treatment groups after transplantation.

#### Subcutaneous xenografts

2 x 10^6^ U87 cells expressing either a control or a Rad18-targeting shRNA were injected subcutaneously into the flanks on BALB/c nu/nu (nude) in a 1:1 mixture of Matrigel and DMEM at a final volume of 100μL (5 mice/group). Mice were treated with doxycycline (Sigma, 2 mg/mL) in drinking water every 48 hours. Tumor growth [(length x width x thickness)/2] was measured using calipers.

#### Orthotopic xenografts

Orthotopic xenografts were established with primary glioblastoma cells expressing a stable shRNA (Gb4) in 4-6 weeks-old female nude mice (Janvier Labs, France). Mice received a stereotaxically guided injection of 2.5 x 10^5^ cells resuspended in 5 µL of DMEM-F12. The injection was precisely located into right forebrain (2 mm lateral and 1 mm anterior to bregma at a 5 mm depth from the skull surface). Survival curves were established and mice were sacrificed at the appearance of neurological signs. Excised brains were collected for subsequent pathology and immunohistochemistry analysis.

### Immunohistochemistry

Immunohistochemical (IHC) stain was performed using the AS48 autostainer platform (Agilent) on FFPE tissue sections (3 µm). After dewaxing, tissue slides were heat pre-treated using EnVision FLEX target retrieval solution Low pH (pH6) for 20 minutes at 97 °C. The slides were blocked for endogenous peroxidase activity and incubated with primary anti-RAD18 antibody (1/500, Envision FLEX diluent, ab188235, Abcam).

The target is visualized using Envision FLEX reagent HRP and DAB, included in the kit (K80021, Agilent). Slides were counterstained using Hematoxylin, dehydrated and mounted using a xylene-based mounting medium.

### Immunohistochemistry scoring

The intensity of RAD18 immunochemistry is evaluated by the H score and the ImmunoReactive Score (IRS). The H-score is a method of assessing the extent of nuclear immunoreactivity. The score is obtained by the formula: 3 x percentage of strongly staining nuclei + 2 x percentage of moderately staining nuclei + percentage of weakly staining nuclei, giving a range of 0 to 300. The IRS system (IRS) was a combination of the intensity (0 to 3 points) and proportion of positive cells (1: < 10 %; 2: 10-50 %; 3: 50-80 %; 4: > 80 %), giving a range of 0 to 12. RAD18 was considered as positive when the H score was > 80 and IRS was > 6.

### Statistical analysis

A linear mixed-regression model, containing both fixed and random effects, was used to determine the relationship between tumor growth and days after grafting. Data were first transformed using the natural log scale to better fit the assumptions of the linear mixed model. The fixed part of the model included variables corresponding to the number of postgraft days and the different treatments. Interaction terms were built into the model; random intercepts and random slopes were included to take time into account. The model coefficients were estimated by maximum likelihood and considered significant at the 0.05 level. Statistical analysis was performed on GraphPad Prism 5.

## Results

### Rad18 is required for GBM cells proliferation

By performing a more detailed analysis of phenotypes associated with Rad18 downregulation in GBM, we have observed a strongly reduced proliferation capacity in four different glioblastoma adherent cell lines treated with either an shRNA (Figure 1A-B), or a previously validated siRNA of different sequence targeting Rad18 (Figure 1C), (6), (Supplementary Figure 1A, and see Materials and Methods). Importantly, these phenotypes were not observed upon Rad18 downregulation with the same siRNA in either HEK293T cells, or in a multiple myeloma cell line (Supplementary Figure 1B-C), consistent with the notion that Rad18 is not essential for life (12–16).

**Figure 1.**
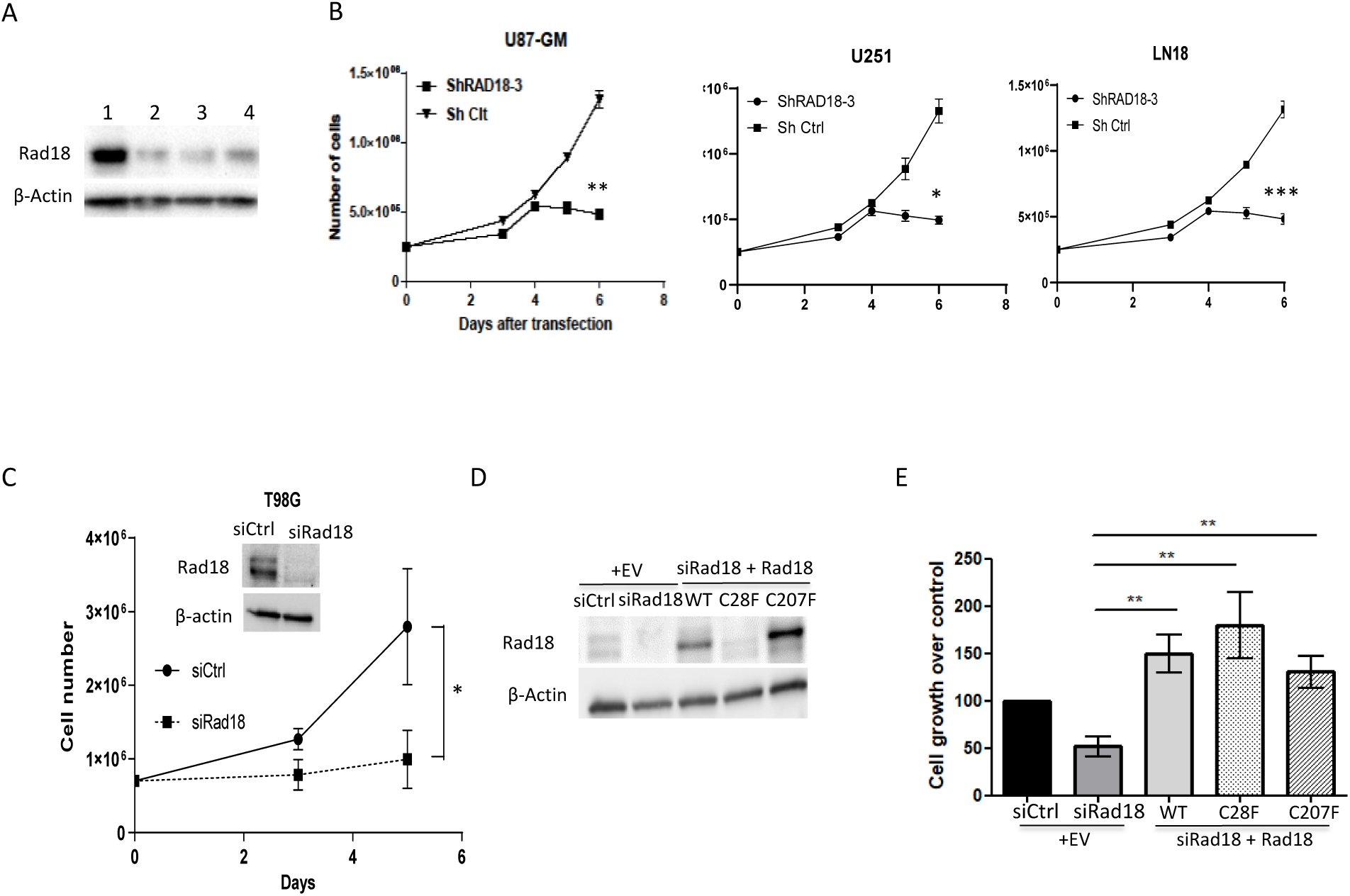
Rad18 downregulation inhibits GBM cells growth. (A) Western blot total protein extracts prepared from U87 (2), U251 (3) LN18 (4) GBM cells treated with either control (lane 1) or Rad18 shRNA (lanes 2-4). n=3. (B) Proliferation of the indicated cells treated with either control (Ctrl) or Rad18 shRNA. Data are means ± SD. *p<0.05; **p<0.01; ***p<0.001. n= 3. (C) Proliferation of T98G cells treated with either control (Ctrl) or Rad18 siRNA. Data are means ± SD. *p<0.05. Inset, western blot of total T98G cell extracts treated with the indicated siRNA. n=4. (D) Western blot of total T98G cell extracts treated with the indicated siRNA and transfected with either empty vector (+EV) or vectors expressing the indicated Rad18 forms (siRad18 + Rad18). n=5 (E) Quantification of T98G cells growth treated as indicated in (C). n= 5. Data are means ± SD. **p<0.0025. Each bar of the graph was normalized as follows: 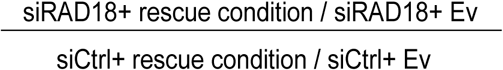

The proliferation defect we observed is much stronger that the one previously reported in GBM adherent cell lines (8) which we believe is due to a hypomorphic effect due to incomplete Rad18 downregulation. Notwithstanding, the specificity of the proliferation defect associated with Rad18 downregulation has not been previously assessed. Hence, we complemented GBM cells treated with siRNA targeting Rad18 with either Rad18 wild-type, or two different mutants targeting two distinct Rad18 functions, its ubiquitin ligase domain (Ring-finger, mutant C28F, involved in TLS) or its zinc-finger domain (C207F mutant, implicated in DSBs repair). We observed that the proliferation defect was completely rescued by expression of either Rad18 wild-type or the two mutants (Figure 1E). This result excludes potential off-target effects of the siRNA but also shows that the proliferation function of Rad18 is independent of its catalytic activity. Interestingly, in conjunction with a strong proliferation defect, cells also displayed a marked change in morphology, presenting clear linear protrusions (Supplementary Figure 1B). Altogether, these results show that Rad18 expression is key to proliferation of GBM adherent cell lines.

### Rad18 downregulation in GBM leads to cell cycle arrest in the G1 phase of the cell cycle

To better understand the consequences of Rad18 downregulation on GBM proliferation, we analyzed the cell cycle profile by flow cytometry, in the presence of the nucleotide analogue BrdU. Results show that Rad18 downregulation by siRNA resulted in a both strong inhibition of BrdU incorporation and accumulation of cells in the G1 phase of the cell cycle compared to cells treated with a control siRNA (Figure 2A-B). Because Rad18 is involved in relieving replication stress, as well as in DSBs repair, the G1 arrest could be explained as a consequence of accumulation of DNA damage leading to cell cycle arrest mediated by the DNA damage response. However, we did not observe an increase in the level of p53, nor its activation by phosphorylation, although we observed an increase in the level of the cyclin-dependent kinase inhibitor p21 (Figure 2C). We also did not observe an increase in the level of the non-specific DNA damage marker γH2AX (Figure 2D, Supplementary Figure 1E), nor that of the phosphorylated form of the Chk1 kinase, a replication stress marker, suggesting that the G1 arrest may not be mediated by the DNA damage response. Finally, we did not observe changes in the total level of its TLS partner Rad6 (Figure 2D). Altogether, these results show that Rad18 downregulation in GBM adherent cells leads to cell cycle arrest in G1 in the apparent absence of DNA damage.

**Figure 2.**
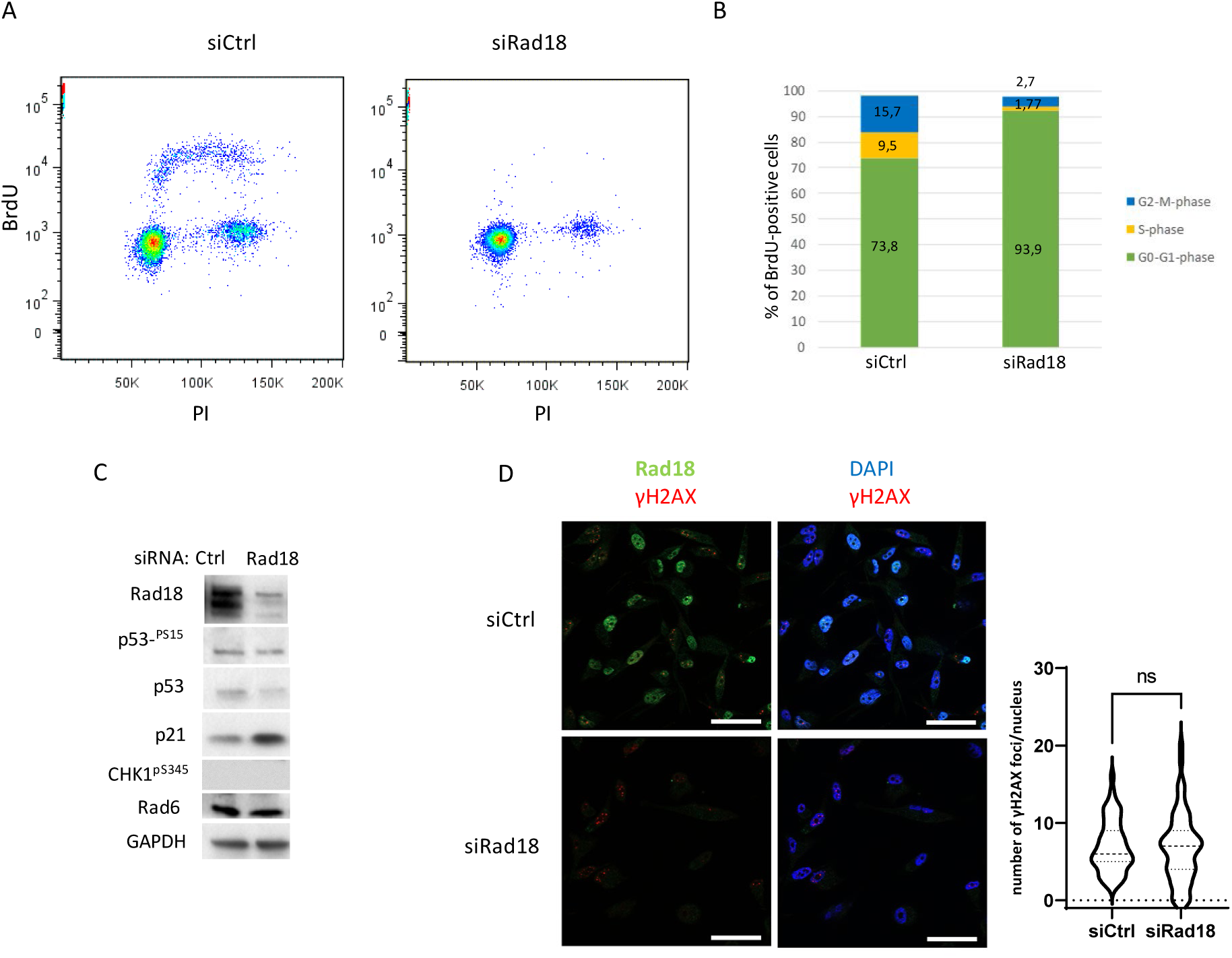
Rad18 downregulation arrests GBM cells in the G1 phase of the cell cycle. (A) BrdU incorporation assessed by flow cytometry of U87 cells treated with either control (Ctrl) or Rad18-specific siRNA. Data were plotted against the intensity of propidium iodide (PI) as a measure of DNA content. n=3. (B) Quantification of cells in the different phases of the cell cycle of the data shown in (A). (C) Western blot with the indicated antibodies of total protein extracts prepared from U87 cells treated with either control (Ctrl) or Rad18 siRNA. (D, left panel) Detection of the indicated proteins by indirect immunofluorescence with specific antibodies. DAPI was used to stain nuclei. Scale bar= 40 µm. Right panel, quantification of the immunofluorescence images shown in the left panel. ** p<0.0017, non-parametric Mann Whitney test. ns, non-significant. n=3.

### Rad18 sustains GSCs stemness

We next explore whether Rad18 may also be important for the proliferation of patients-derived primary tumors enriched in GSCs. To this end, we treated two different GSCs isolated from primary tumors (Gb4, Gb7)(17) with either control or Rad18-specific siRNA. Rad18 downregulation by siRNA in these cells resulted in a strong reduction of the size of the gliomaspheres (Figure 3A-B). Consistent with this result, Rad18 downregulation decreased the expression level of the cancer stem cell markers Sox2 and Oct4 (Figure 3C), and consistent with results shown in Figure 2C-D, did not increase γH2AX levels. These phenotypes were further confirmed by targeting Rad18 expression with an shRNA that did not share any sequence homology with the abovementioned siRNA (Supplementary Figure 2A-B). To assess a role for Rad18 in GSC frequency and self-renewal, we performed extreme limited dilution assay (ELDA, Figure 3D-E). As can be seen, Rad18 downregulation resulted in marked decreased capacity of GSCs to form spheres, supporting a role for Rad18 in GSC self-renewal.

**Figure 3.**
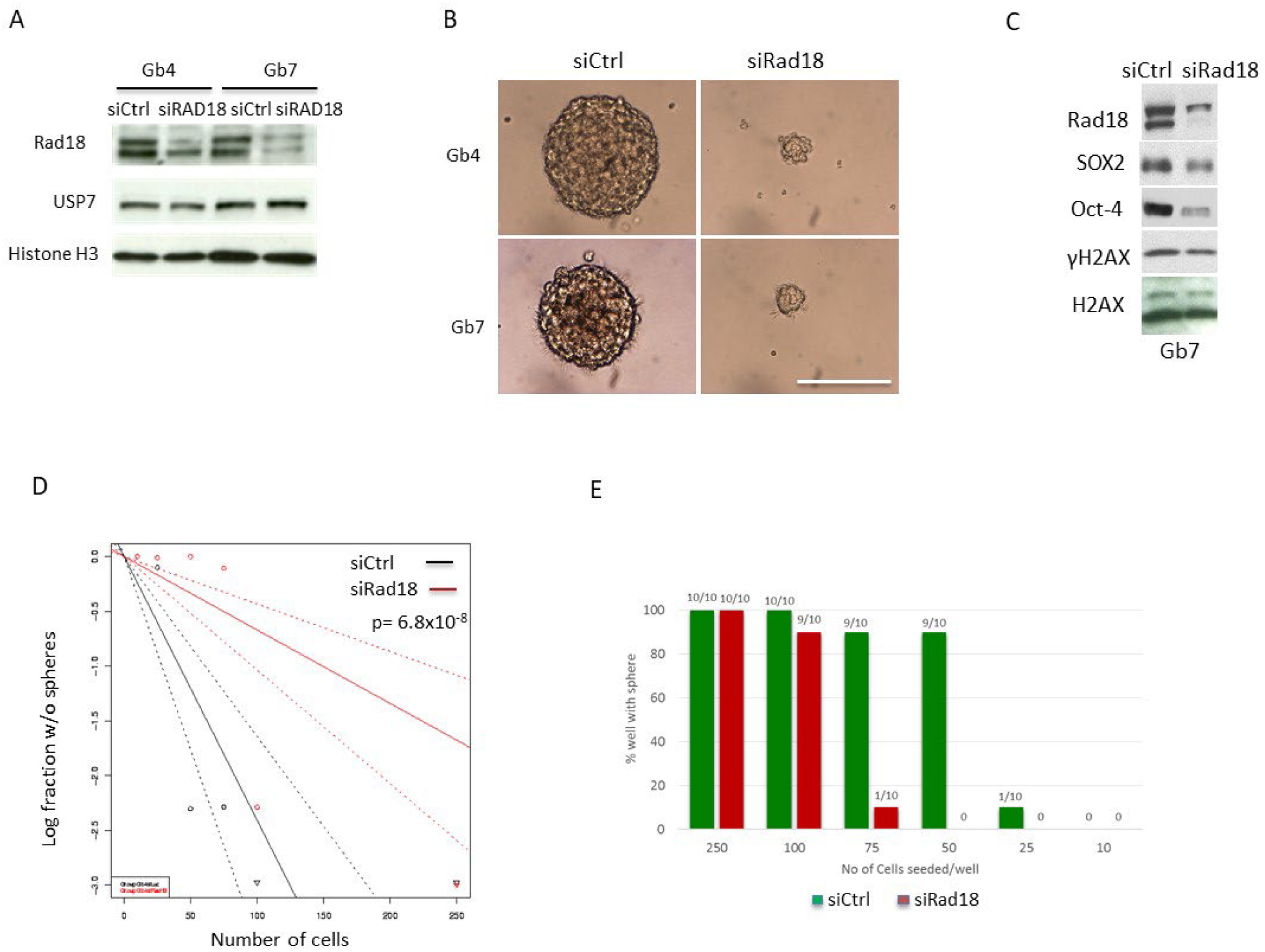
Rad18 downregulation strongly inhibits GSCs growth. (A) Western blot of total cell extracts with the indicated antibodies obtained from either Gb4 or Gb7 GSCs treated with either control (Ctrl) or Rad18 siRNA. n=3 (B) Representative phase contrast images of Gb4 cells treated with either control siRNA (siLuc) or Rad18 siRNA. Scale bar: 100 µm. (C) Western blot of total cell extracts with the indicated antibodies obtained from Gb7 GSCs treated with either control (Ctrl) or Rad18 siRNA. n=3. (D) Limited dilution assay obtained with Gb4 cells treated with either control (black lines) or Rad18 siRNA (red lines) calculated with extreme dilution analysis (ELDA). (E) Number of cells per well required to form gliomaspheres of Gb4 cells treated with either control siRNA (siLuc) or Rad18 siRNA. n=4.

### Rad18 downregulation reduces tumor growth *in vivo*

Although Rad18 has been more recently implicated in the growth of GBM adherent cells in culture (8), its role in glioblastoma growth *in vivo* has not been previously assessed. To this end, we analyzed tumor formation by performing mouse xenografts with transgenic GBM cells expressing an shRNA targeting Rad18 of a different sequence of the abovementioned sh- and siRNAs (see Materials and Methods). First, we tested tumor growth by generating subcutaneous xenografts with adherent GBM cells. Cells were injected under the skin of immunocompromised scid nude mice and tumor growth was analyzed compared to cells expressing a non-specific shRNA (control). While we observed exponential tumor growth in mice grafted with control shRNA as expected, tumor growth obtained with cells expressing Rad18 shRNA was strongly reduced (Supplementary Figure 2C). Quantification of tumor size confirmed a reduction in tumor volume from mice injected with GBM cells expressing the shRNA targeting Rad18 (Supplementary Figure 2D). Consistent with these results, mouse survival was significantly increased (Supplementary Figure 2E). This result was formally confirmed by performing orthotopic xenogratfs in the mice brain with GSCs expressing the same inducible shRNA targeting Rad18 (Supplementary Figure 3A-B). Analysis of brain slices by staining with either hematoxylin & eosin or with a Rad18-specific antibody (Supplementary Figure 3C) shows that GSCs in which Rad18 expression was inhibited, were unable to form a large tumor mass, and appeared to expand without connections between them in the mice brain. (Figure 4A-B)

**Figure 4.**
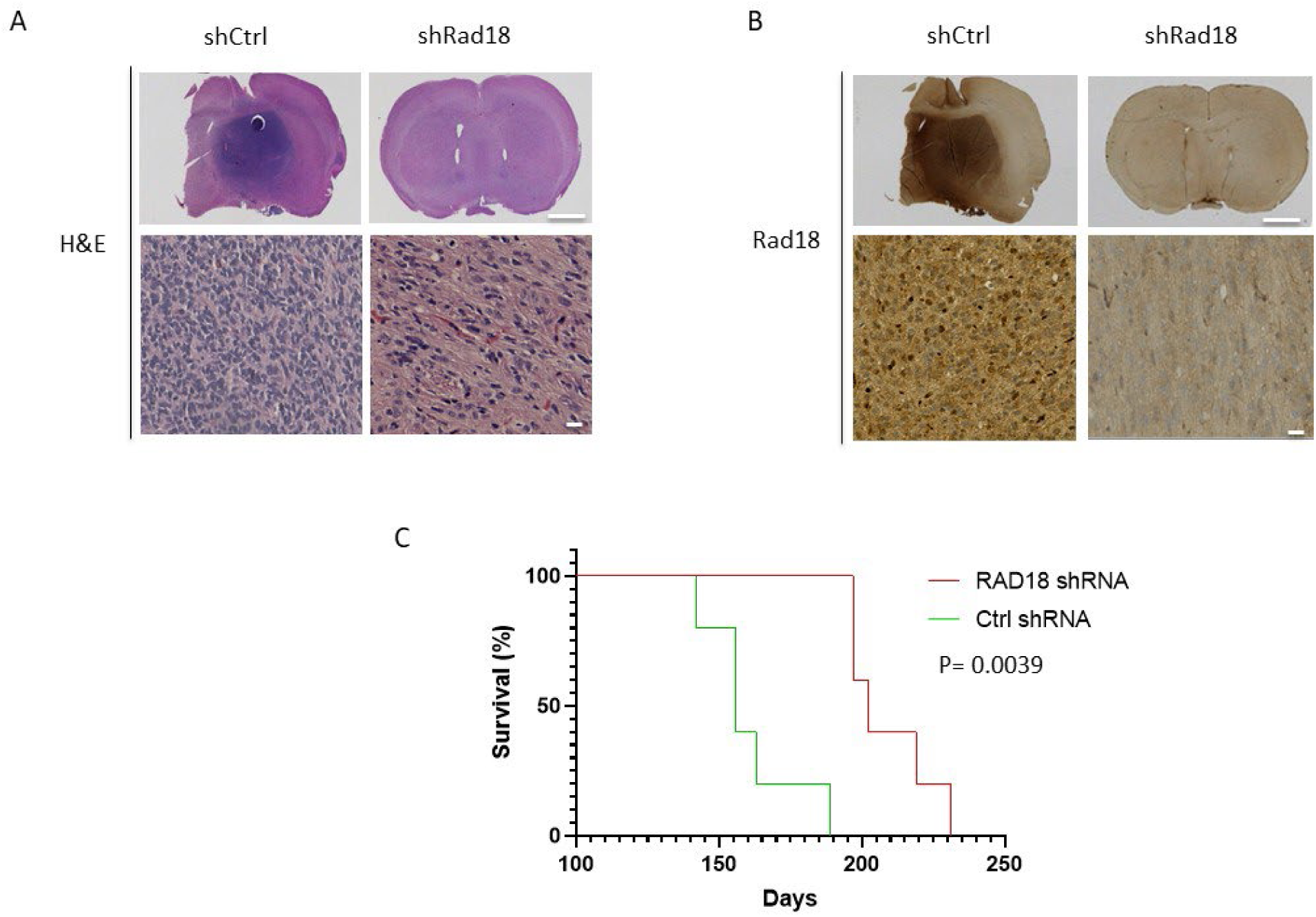
Rad18 downregulation reduces GBM growth in mice orthotopic xenografts. (A) Hematoxilin and Eosin (H&E) staining and (B) Rad18 staining by immunohistochemistry of cross sections of mouse brains bearing GSC-derived tumors from mice injected with GSCs expressing an shRNA control (Ctrl) or and shRNA targeting RAD18. Scale bar: 2 mm. Lower panels show magnification of the upper panel images. Scale bar: 20 µm. (C) Kaplan-Meier survival curves of mice intracranially transplanted with GSCs expressing the indicated shRNA. Log-rank analysis was used. n= 5 mice per group.

Consistent with this result, inhibition of Rad18 expression significantly extended mouse life span (Figure 4C). Altogether, these observations show that inhibition of Rad18 expression strongly reduces the growth of GBM xenografts in mice and demonstrate that Rad18 is essential to sustain GBM growth *in vivo*.

### Analysis of Rad18 expression in GBM specimens

The reason of Rad18 upregulation in GBM is currently unknown. The Rad18 gene is located on chromosome 3p25.3, a locus which is not amplified in glioblastoma (18). Rad18 is a target of the E2F3 transcription factor, a critical regulator of cell cycle progression (19). A previous report showed that in GBM, E2F3 interacts with the chromatin remodeler complex HELLS and that this interaction is important to activate E2F3 target genes (20). Interestingly, HELLS is preferentially expressed in GSCs, similar to Rad18 (6), and its downregulation strongly inhibits GBM proliferation (20). Consistent with this possibility, correlation analysis using the Cancer Genome Atlas for Glioblastoma Multiforme (TCGA-GBM) dataset (Rembrandt and Bao collection) shows a tight correlation between Rad18 and HELLS expression, and not between Rad18 and c-myc, this latter being another HELLS target (Supplementary Figure 3D-F). Further, no positive correlation between the expression of Rad18 and PDGFRA could be observed (Supplementary Figure 3G), in contrast to a previous report showing increased PDGFRA expression upon Rad18 knockdown (8), and consistent with the notion that high PDGFRA expression in linked to a poor prognosis (21). Taken together, these observations suggest that high expression of Rad18 in GSCs may be a direct consequence of high HELLS expression.

Data mining of Rad18 expression in the TCGA-GBM dataset also revealed that Rad18 is upregulated in GBM compared to normal tissues (Supplementary Figure 4A, left panel), and that high Rad18 expression correlates with poor overall survival (Supplementary Figure 4A, right panel). These observations corroborate results obtained in mice xenografts (Supplementary Figure 2E and Figure 4), and indicate that Rad18 expression has prognostic value, in line with two recent reports (7, 9). Rad18 expression increases with the GBM grade (Supplementary Figure S4B-C), while no significant differences were observed between GBM classes (Supplementary Figure 4D). Further, Rad18 expression is higher in GBM having a wild-type IDH1 status, which is an indicator of poor prognosis (Supplementary Figure 4E). Comparison of Rad18 expression in GBM with that of RNF8 and RNF168, two other E3 ubiquitin ligases involved in similar pathways to Rad18, shows that these latter are rather underexpressed in GBM (Supplementary Figure 4F-G) and that their prognostic value is much lower than that of Rad18. Further, amongst three Y-family TLS pols, functionally linked to Rad18, only Polκ was previously shown to serve as an independent prognostic factor in glioma in a cohort of 104 patients (22). However, comparison of Rad18 expression with that of Polκ in the TCGA database, shows that Rad18 expression is higher than that of Polκ and that Rad18 has better prognostic value than Polκ (Supplementary Figure S4A, H). Altogether, these observations support the experimental data and are indicative of a specific, TLS-independent functional role for Rad18 in GBM progression.

### Increased proliferation potential and loss of contact inhibition upon ectopic Rad18 expression in mammalian cells

Analysis of GBM patients’ specimens shows that biopsies having a high proliferation MIB index also have an increase of Rad18 expression (Figure 5A). Staining of GBM biopsies specimen with a Rad18-specific antibody (Supplementary Figure S3C) showed two different stains: a strong stain in 1 to 80% of nucleus cells and in a light non-specific stain of the cytoplasm in GBM cells and in the fibrillary background (Figure 5B). This observation suggested an increased proliferation potential of RAD18. The strong nuclear stain was in the line with the observation that Rad18 is upregulated in GSCs (6).

**Figure 5.**
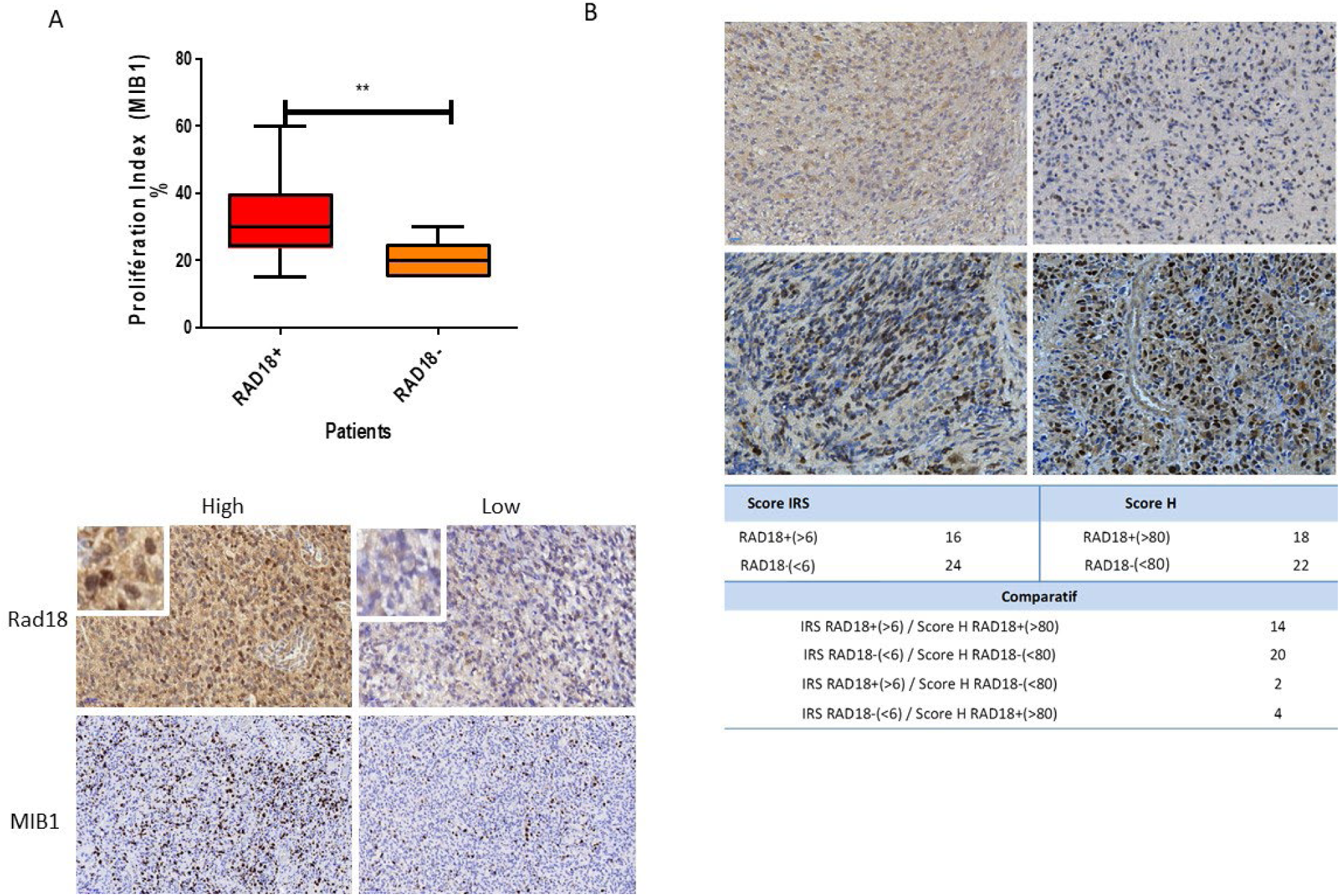
Rad18 expression in human GBM specimens. (A, upper panel), proliferation index (MIB1) determined in human GBM grade IV biopsies issued from patients having high (Rad18+) or low (Rad18-) Rad18 expression. Lower panel, immunohistochemistry staining of grade IV GBMs biopsies with either Rad18 or MIB1, presenting a high or low reactivity. Insets show magnifications of part of the panels. **P<0.0028 (unpaired t-student test). Scale bar: 50 µm. (B, upper panel) examples of Rad18 detection in a set of human grade IV GBM primary tumors by immunohistochemistry. Scale bar 20 µm. Lower panel, nuclear immonoreactivity score (H score) and the ImmunoReactive Score (IRS) of grade IV GBMs stained with Rad18.

These observations suggest that Rad18 may have an intrinsic oncogenic potential. To test this possibility, we analyzed the growth of non-transformed mouse NIH3T3 cells expressing Rad18 in soft agar, a stringent assay to test for anchorage-independent growth, a hallmark of malignant transformation. As can be seen (Figure 6A), NIH3T3 cells expressing ectopic Rad18 have a greater proliferation advantage than control cells expressing the empty vector (Figure 6B). Further, we determined whether in addition to increased anchorage-independent proliferation, these cells might have lost the ability to arrest proliferation once reached confluence. For this, we analyzed the ability of these cells to form colonies revealed by staining with crystal violet. Results shown in Figure 6C show that cells expressing ectopic Rad18 and not the empty vector had the ability to form colonies, suggesting acquisition of oncogenic properties. In line with our previous observations (6), cells expressing ectopic Rad18 at somehow similar levels than endogenous Rad18 suppress activation of the DNA damage checkpoint induced by UV-irradiation. Altogether, these observations show that increased Rad18 expression has oncogenic potential and is able to override the DNA damage response, thus facilitating cell proliferation.

**Figure 6.**
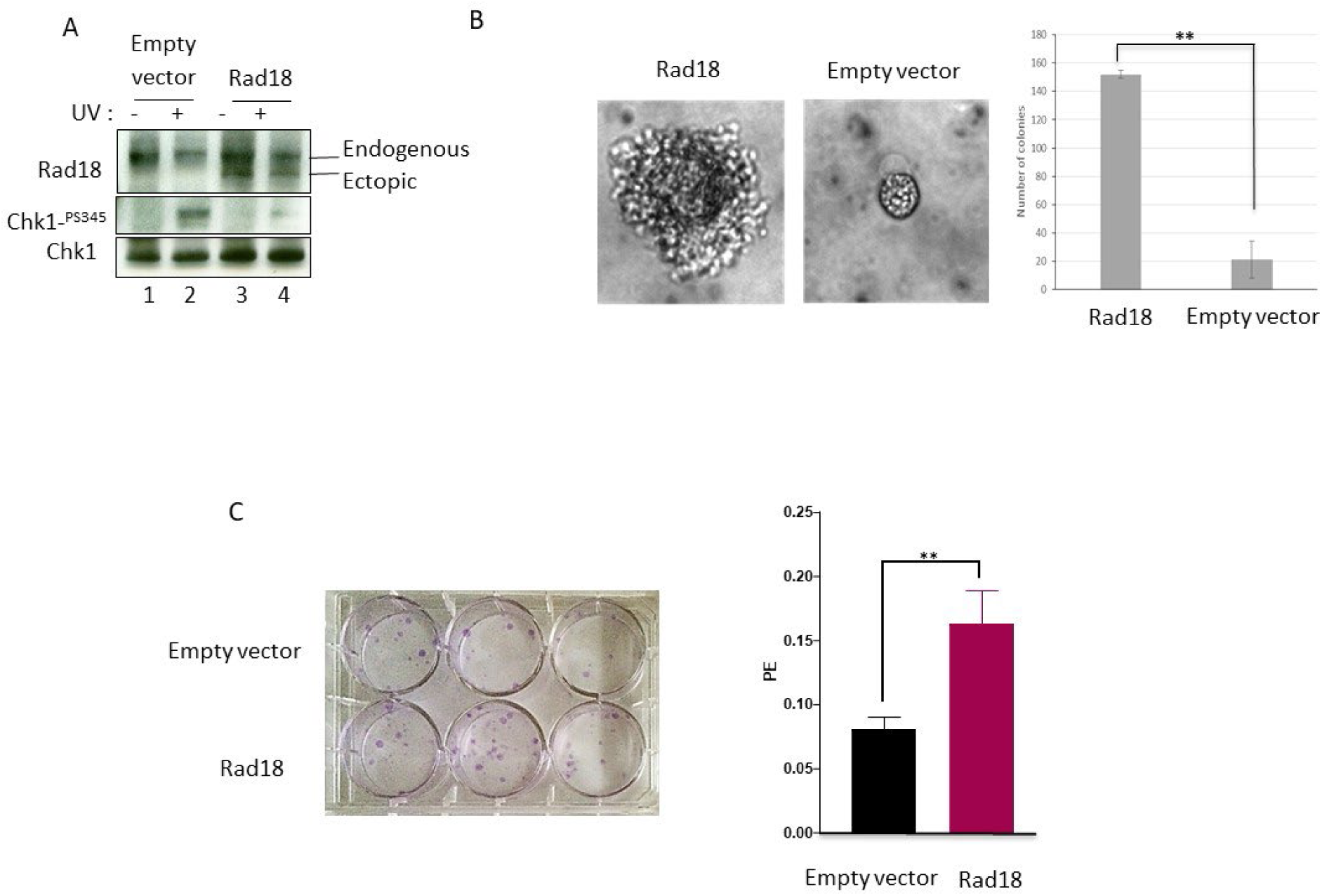
Ectopic Rad18 expression stimulates growth of mouse NIH 3T3 cells. (A) Western blot with the indicated antibodies of total cell extracts obtained from stable NIH 3T3 cells expressing either empty vector (pCDNA3) or ectopic Rad18. n=3. (B, left panel) Representative phase contrast images of NIH 3T3 cells expressing either empty vector or Rad18 growing in soft agar. Right panel, quantification of data shown in left panel. n=5. (C, left panel) Colony formation assay of stable NIH 3T3 cells described in (A) expressing either empty vector or Rad18. Right panel, quantification of the data shown in the left panel. Data are expressed as plating efficiency (PE) and are means ± SD. ** p<0.0059. n=3.

### Rad18 controls the Hippo pathway in GBM

A very recent report has suggested that Rad18 stimulates the Hippo-YAP pathway in triple negative breast cancer cells (23). Because as Rad18, this pathway is upregulated in GBM (24), we investigated its possible functional relationship with Rad18. Interestingly, we observed that upon Rad18 downregulation, YAP levels were decreased, and a slow migrating polypeptide recognized by the YAP antibody was also detected, suggesting phosphorylation (Figure 7A). In contrast, the level of other factors involved in this pathway were not significantly changed (Figure 7B-C). YAP phosphorylation occurs in the cytoplasm leading to its degradation by the proteasome. Consistent with this notion, analysis of YAP localization by indirect immunofluorescence shows that upon Rad18 downregulation, cytoplasmic YAP is increased compared to control samples (Figure 7D). Finally, if reduced YAP activity accounts for the proliferation slow down observed upon Rad18 depletion, it is expected that YAP expression should rescue the proliferation defect of Rad18-depleted cells. As can be seen in Figure 7E, YAP rescued the proliferation defect of Rad18-depleted cells. Altogether, these results show that Rad18 is involved in the negative regulation of the Hyppo pathway.

**Figure 7.**
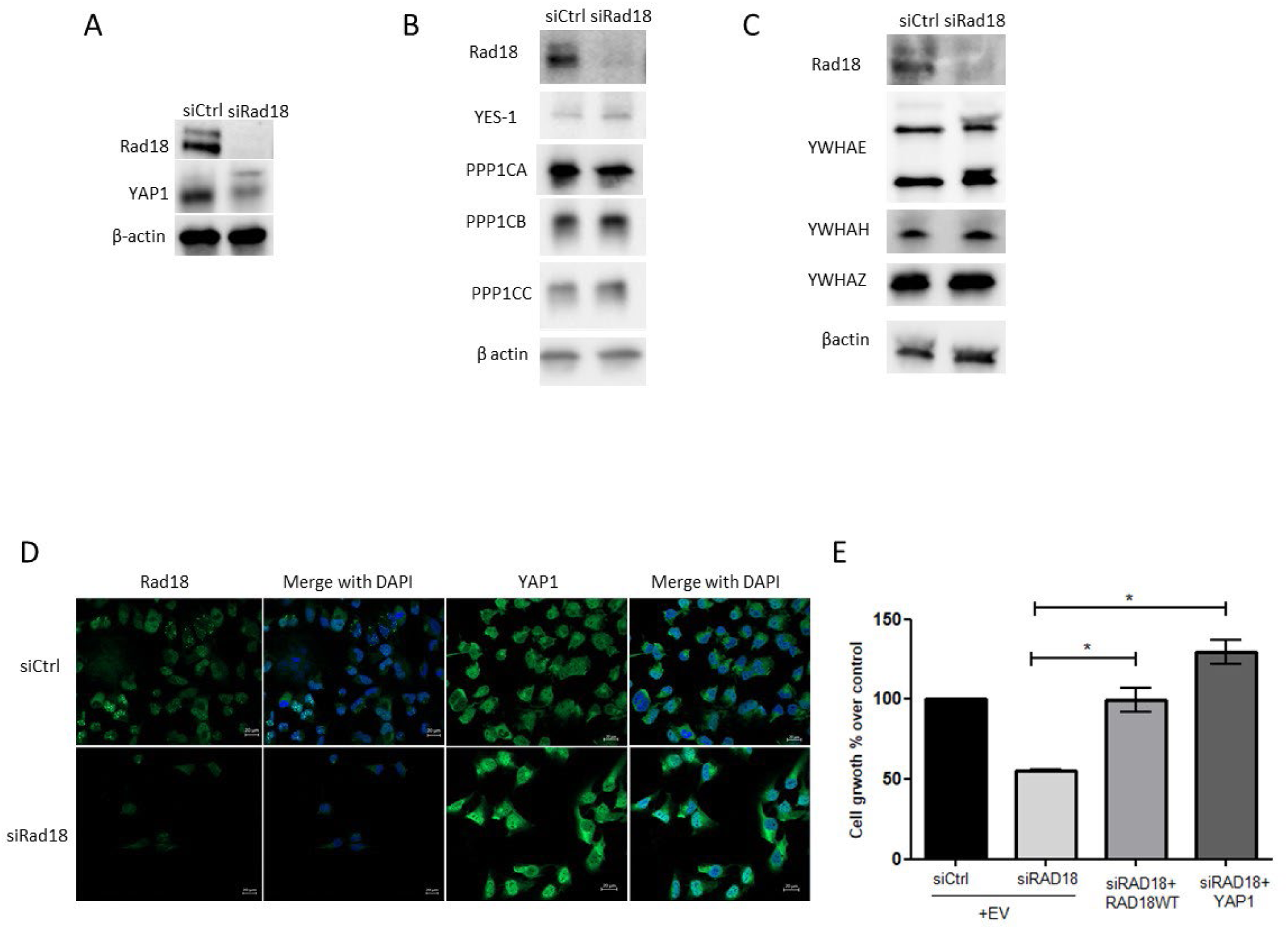
Rad18 regulates the Hyppo pathway. (A-C) Western blot of total cell lysates obtained from either T98G (A) or GCSCs (B) upon transfection with the indicated siRNAs. n=3. (D) Immunofluorescence images of T98G cells treated with the indicated siRNAs. DAPI was used to visualize DNA. n=2. (E) T98G cells viability upon transfection with the indicated siRNAs and the indicated expression vectors. EV: empty vector. * P=0, 028 siRad18+Rad18 WT; P=0, 01 siRad18+YAP1. n=2.

## Discussion

Rad18 is a non-essential gene in yeast and vertebrate cells. Rad18 knockout mice develop normally, although males are partially sterile (13). Rad18 downregulation was previously reported not to affect proliferation of either normal, or cancer cells, neither that of embryonic stem cells, but sensitizes them to a wide range of DNA damaging agents (14, 25–28), as expected given its known function in DNA damage tolerance and DSBs repair. Notwithstanding, recent studies have implicated Rad18 in tumorigenesis, in part by facilitating tolerance to oncogenic stress (29), and also by promoting mutagenesis (30).

We (6) and others (31) have previously shown that Rad18 is overexpressed in GBM, and two recent reports have implicated Rad18 in the proliferation of GBM adherent cell lines, although the specific role of Rad18 in this process has not been investigated (8, 32). In this report, we have shown that Rad18 is surprisingly essential for GBM cells proliferation, as well as for GBM progression in mouse preclinical models, in the absence of external DNA damage. The reduced proliferation capacity of GBM adherent cells we have observed is consistent with two previous reports (8, 32), although we observed a much stronger effect, that we believe is due to a higher depletion of Rad18 in our experiments.

We have also shown that the proliferation arrest is a result of a strong delay in the G1 phase of the cell cycle, which may explain a previous observation showing reduced level of the G1/S transition regulator cyclin D1 (32). The cell cycle arrest appears to be independent of the status of the tumor suppressor p53, since it was observed in both p53 wild-type (U87 an dU251) and p53 mutated cell lines (LN18 and T98G). We have also shown that Rad18 downregulation strongly reduces the self-renewal capacity of GSCs and reduces the expression level of GSCs markers, such as Sox2 and Oct4. Interestingly, the proliferation defect observed by Rad18 downregulation in GBM in the absence of external damage was not observed upon downregulation of TLS Polκ (33), suggesting that Rad18 may have a TLS-independent function in GBM proliferation. Because TLS is mutagenic, this possibility is consistent with the notion that GBM has a low mutation load, while this strongly increased tremendously upon therapeutic treatment (34). Consistent with this scenario, TLS Polκ downregulation was reported to strongly increase the life span of GBM mice xenografts exposed to temozolomide (33).

Further, we have observed that Rad18 expression can provide a proliferative advantage in NIH 3T3 transformed cells, and that the proliferation defect observed upon Rad18 downregulation in GBM is independent of its known catalytic activity. This scenario also explains why Rad18 downregulation does not apparently induce DNA damage in GBM cells, in contrast to a previous report showing that Rad18 downregulation in GBM cells increased the level of the DNA damage responsive gene p53 (32). This difference may be due to a difference in the cell lines used and/or to hypomorphic effects as a result to partial Rad18 depletion.

In this report, we have also provided new insights into how Rad18 functions in promoting GBM growth. Consistent with a very recent report in breast cancer cells (23), we have observed that also in GBM, Rad18 may regulate the activation of the YAP-Hippo pathway, though regulation of YAP stability and/or subcellular localization. This result is in line with the observation that YAP upregulation overcomes contact inhibition (35), as observed by Rad18 upregulation in non-transformed cells. Rad18 may regulate either the nuclear or cytoplasmic retention of YAP by a mechanism that remains to be elucidated, although in a catalytically-independent manner.

In conclusion, our observations unveil an important role for Rad18 in sustaining GBM proliferation, and in particular self-renewal of GSCs, and propose Rad18 as a potential target to limit GBM proliferation. Because Rad18 is also involved in DNA damage tolerance and DNA repair, it constitutes an excellent target to improve current GBM therapy, in combination with YAP inhibitors.

## Data Availability

Data supporting the findings of this study are available from the corresponding authors upon request.

## Authors contribution

Conceptualization, D.M. Methodology, D.M., A.S., J.P. Investigation, N.B., C.K., A.S., L.S., J-M.P., J.M., M-B.D., T.E. Writing – original Draft, D.M. Writing – Review & Editing, N.B., C.K., A.S., L.S., J.P., J-P.H., M-B.D., E.U-C., D.M. Funding Acquisition, D.M. Resources, D.M., J-P. H., E.M., E.U-C. Supervision, D.M., E.U-C., J.P, M-B.D.

## Acknowledgements

This work was supported by grants from ARC (“PROJETS FONDATION ARC” PGA120150202287 to D.M.), Ligue contre le Cancer (to C.K.), MSD Avenir (GNOSTIC to D.M.), GEFLUC (to D.M.), LABEX EPIGENMED (to L. S.) and World Wide Cancer Research (to D.M. and N. B.). We thank T. Virolle for critical reading of the manuscript. The authors declare that there are no financial conflict of interests that might be construed to influence the results or the interpretation of this manuscript.

## Supplementary Information

### Includes four Supplementary figures

**Figure S1.**
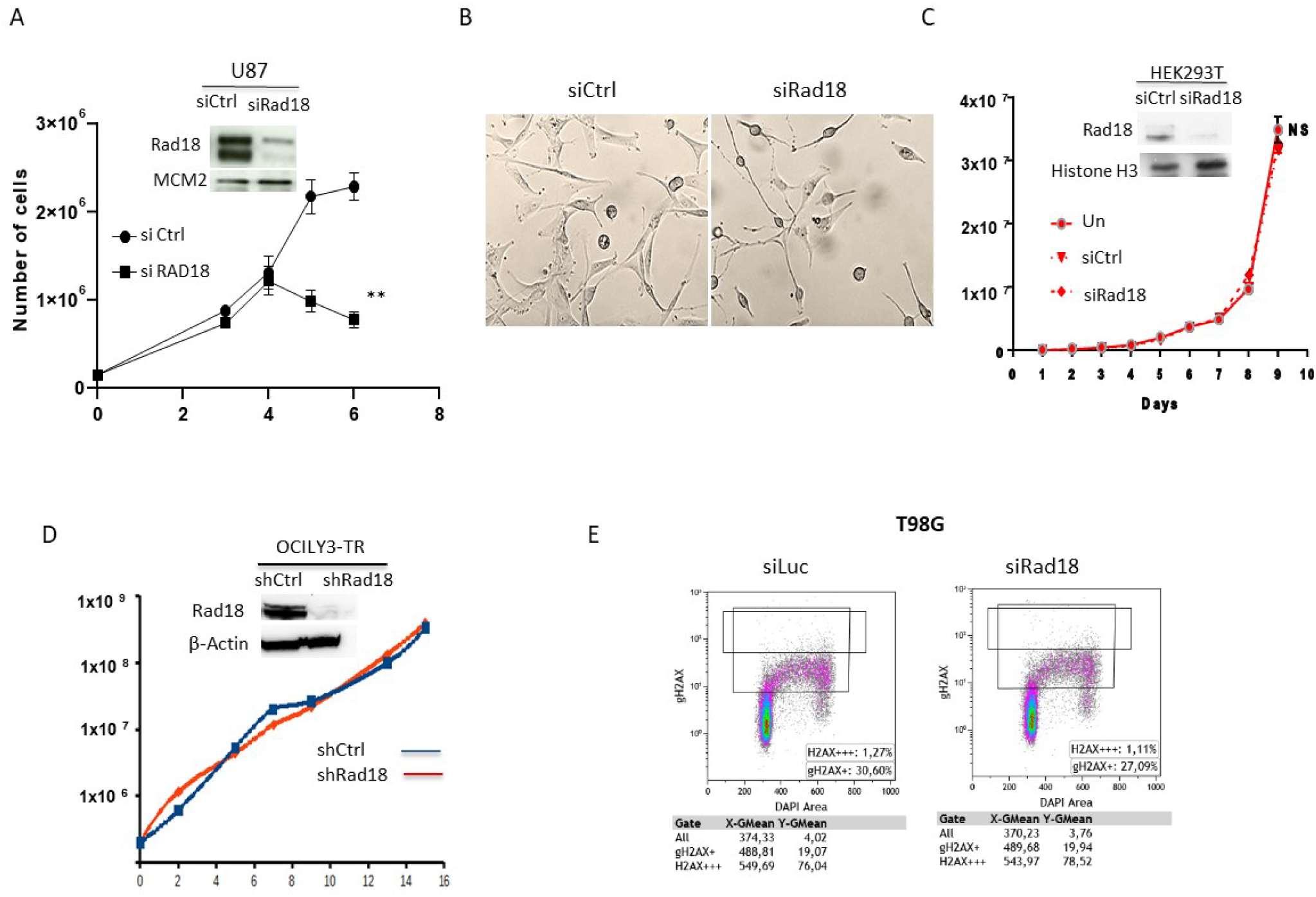
(A-D) Proliferation of the indicated cell lines treated with either control (Ctrl) or Rad18 siRNA (A-D). Insets show western blots of total cell extracts obtained from the indicated cells to show the efficiency of Rad18 depletion. (B) phase contrast photographs of U87 cells treated as indicated at the 96 h time point. n=3; (E) Detection of γH2AX by flow cytometry in T98G cells.

**Figure S2.**
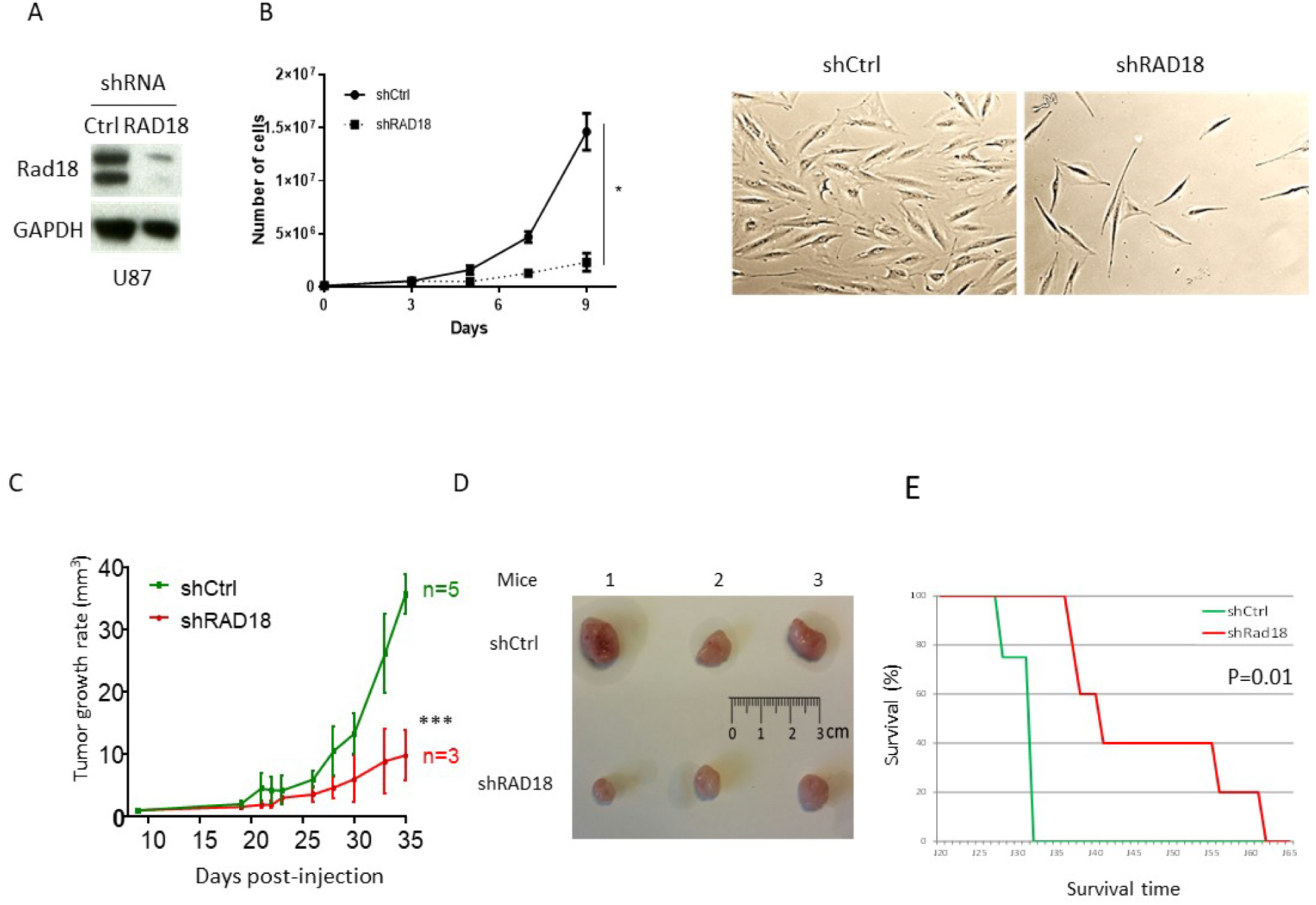
(A) Western blot and (B) growth curve of U87 cells expressing either an shRNA control (shctrl) or an shRNA targeting Rad18. Data are means ± SD. *p<0, 0238. Right panel of Figure S2B shows photographs of cells at the 96 hours time point. (C) Growth rate of subcutaneous xenografts obtained in nude mice subcutaneously injected with U87 cells expressing either an shRNA control (shCtrl) or an shRNA targeting Rad18. Data are means ± SD. ***p<0.0006. (D) Images of tumors obtained in mice treated as described in (C). (E) Kaplan-Meier survival curves of mice treated as in (C). Log-rank analysis was used to calculate survival.

**Figure S3.**
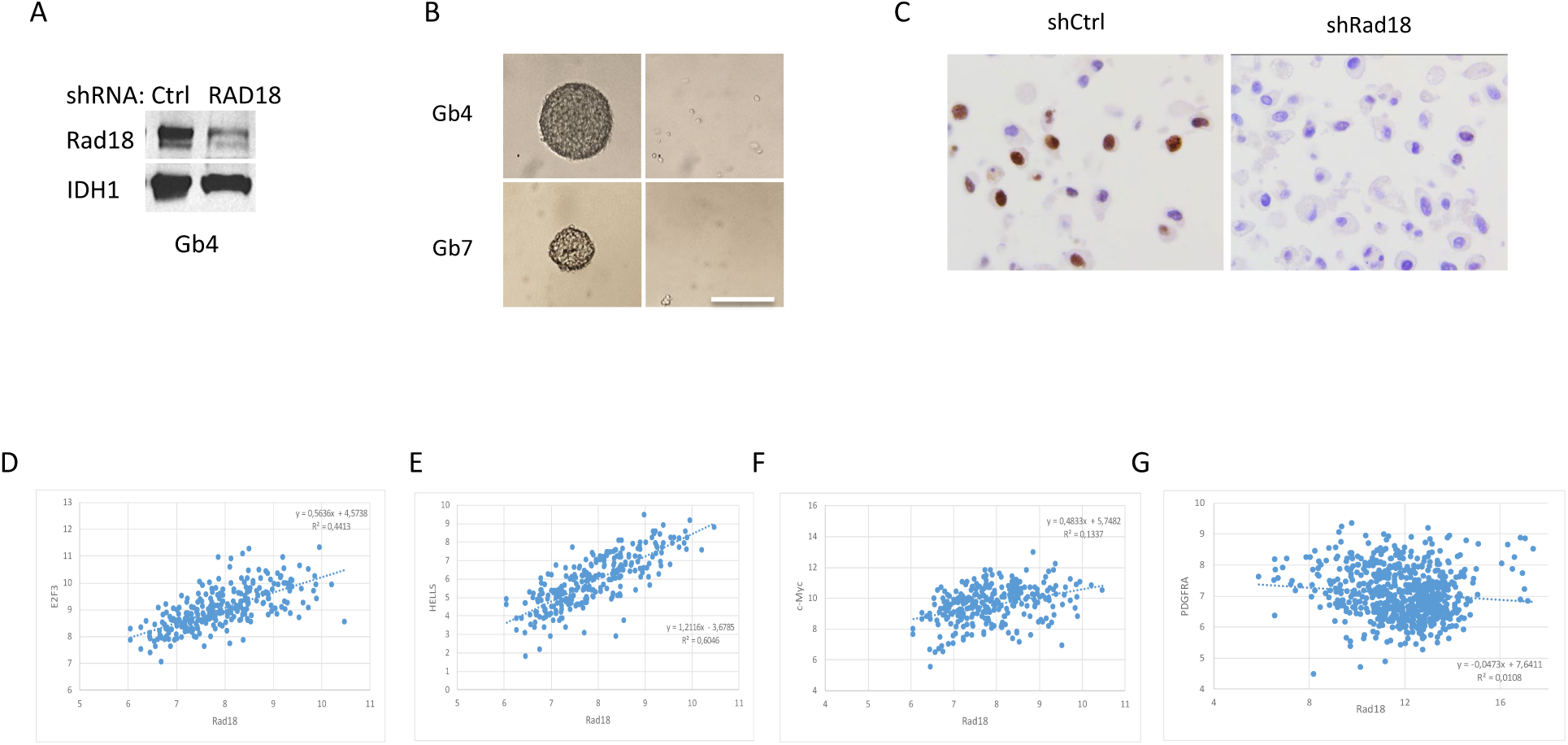
(A) Western blot and (B) phase contrast photographs of Gb4 and Gb7 GCSCs expressing the indicated shRNA. Scale bar: 100µm. (C) Detection of Rad18 by immohistochemistry in U87 cells treated with the indicated shRNA embedded in paraffin. (D-G) Correlation of Rad18 mRNA expression with the indicated genes in GBM obtained from the TCGA.

**Figure S4.**
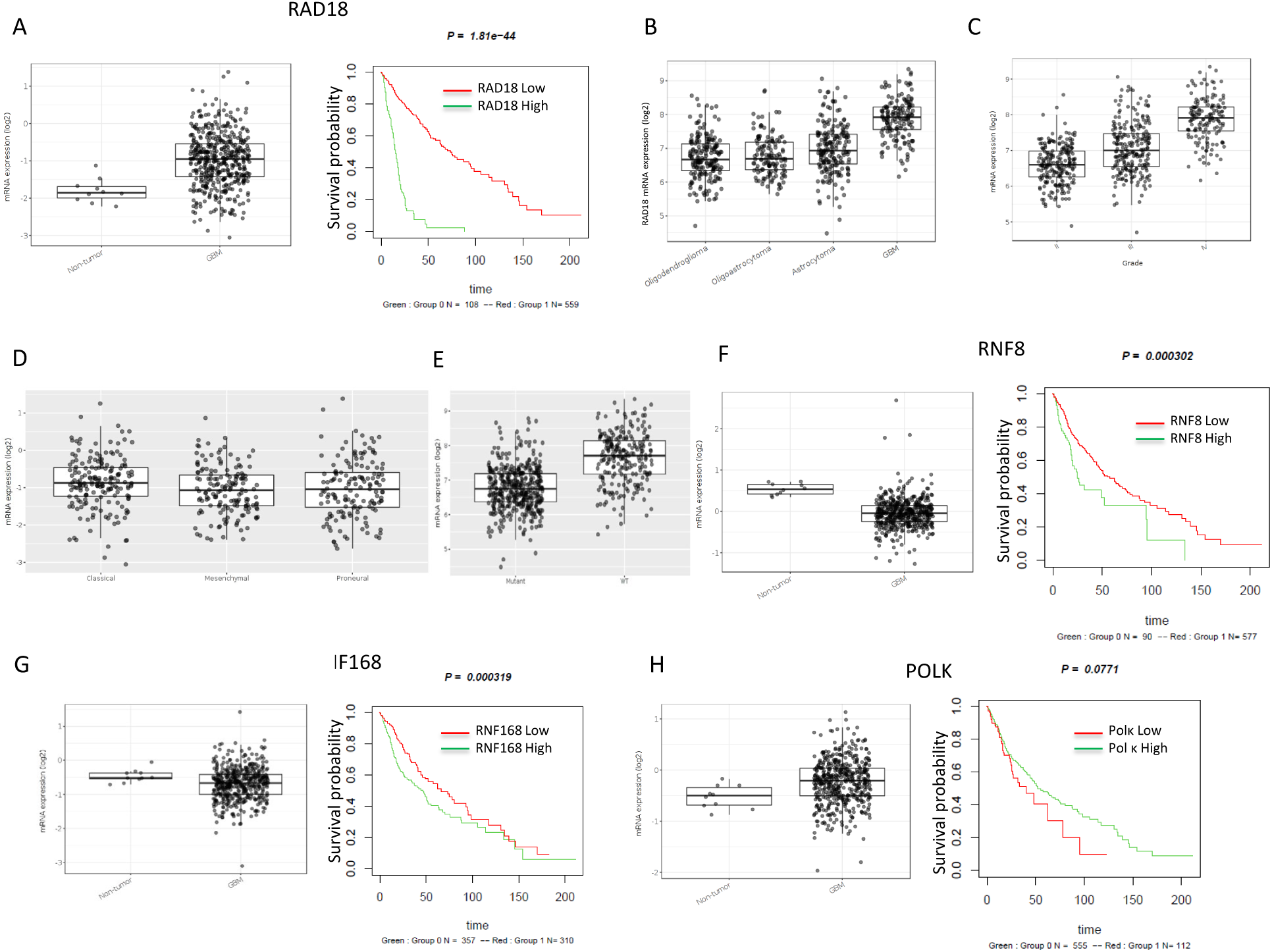
(A-D) Comparison of the indicated genes mRNA expression levels (left panels) and survival (right panels) in normal brain and GBM in the TCGA dataset. (E-H) correlation analysis of the indicated genes mRNA expression on TCGA GBM specimens (Rembrandt and Bao collection). Red line corresponds to high expression, green line to low expression.

